# CasFinder: Flexible algorithm for identifying specific Cas9 targets in genomes

**DOI:** 10.1101/005074

**Authors:** John Aach, Prashant Mali, George M. Church

## Abstract

CRISPR/Cas9 systems enable many molecular activities to be efficiently directed *in vivo* to user-specifiable DNA sequences of interest, including generation of dsDNA cuts and nicks, transcriptional activation and repression, and fluorescence. CRISPR targeting relies on base pairing of short RNA transcripts with their target DNA sequences that must also be adjacent to fixed DNA motifs. However, rules for Cas9 targeting specificity are incompletely known. With increasing numbers of Cas9 systems being developed and deployed in more and more organisms, there is now strong need for a flexible and rational method for finding Cas9 sites with low off-targeting potential. We address this through the CasFinder system, which we demonstrate by generating human and mouse exome-wide catalogs of specific sites for three varieties of Cas9 – *S. pyogenes*, *S. thermophilus* (ST1), and *N. meningitidis* – that each target 56-74% of all exons. We also generate reduced sets of up to 3 targets per gene for use in high-throughput Cas9-based gene knockout screens that target 75-80% of all genes.

## Introduction

Recently, type II prokaryotic Clustered Regularly Interspaced Palindromic Repeats (CRISPR) systems [1,2] have been adapted as tools for efficient targeted genome editing, regulation, and labeling in a rapidly expanding set of eukaryotic hosts [3-15]. In these systems, a complex of a Cas9 protein and a pair of RNAs recognizes a DNA target comprising a short stretch of DNA complementary to the 5’ end of one of the RNAs (referred to as the *spacer* region) followed by a Cas9-specific Protospacer Adjacent Motif (PAM). In the most developed CRISPR system based on *S. pyogenes* (SP) Cas9, the two RNAs are often combined into a single guide RNA (sgRNA) whose 5’-most 20bp comprises the targeting region and whose principal PAM is NGG. However, early studies indicated significant potential for Cas9 off-targeting, and accumulating evidence has only complicated the understanding of Cas9 specificity. While it was recognized initially that matching across the entire 20nt region was not required for Cas9 activity, it was originally thought that mismatches with the last 12-13nt (known as the “seed” region) would abrogate binding [1]. Now it is known both that 1-2 bp seed mismatches can be tolerated and that that base pairing in the region 5’ of the seed also determines activity [7,16-19]. Cas9 recognition of secondary PAM sequences has also been observed [5,20], and recently characterized Cas9 proteins from *N. meningitis* (NM) and *S. thermophilus* (ST1) were found to have unexpectedly complex sets of primary and secondary PAMs [10]. One result of these findings has been to spur efforts to directly engineer Cas9 and its applications to improve specificity, such as the development of paired Cas9-nickases [7,12] and the use of truncated sgRNAs [21]. In the meantime, researchers interested in using Cas9 within genomes of interest have need for better methods of identifying the target sequences that have the lowest potential for off-targets. Recent work with sgRNA libraries has resulted in new algorithms for evaluating Cas9 specificity and corresponding exome-wide catalogs of targets, but these do not cover all targets of potential interest, have considered only SP Cas9, and have not been accompanied by software that other researchers can use [22-24].

## Design and Implementation

The CasFinder system extends and modifies a method we originally developed in ref. [3] of searching for potential Cas9 off-targets using queries that combined seeds and PAMs. The modifications take into account the tolerance to seed mismatches, importance of non-seed matches, and the complexity of PAMs that are now known to apply. These features increase the computational demands on the algorithm and the high variability between the Cas9s of different organisms makes global optimization difficult. Thus, in our present method, we ceded global optimality in favor of developing a simple, practical, and highly flexible system capable of being easily tuned and applied to new genomes and Cas9s. (See Supporting Information S1 for details). Key features include the following:

1. The system implements two distinct functions: (i) finding candidate Cas9 sites within user-specified sequences, performed by the CasFinder program, and (ii) evaluating candidate target sequences in a genome, performed by CasValue. CasFinder calls CasValue to evaluate the candidates it finds, but CasValue may be called independently and can be used to evaluate the specificity of Cas9 targets found by other systems.
2. To evaluate candidate Cas9 targets found by CasFinder, CasValue uses the bowtie [25] sequence mapping tool to retrieve from the user-specified genome all sequences matching the candidate’s seed region up to a specified number of mismatches, and then extends these to full length Cas9 footprints surrounding the matching regions. Footprints whose PAM positions do not contain primary or secondary PAMs for the Cas9 are ignored, and scores are computed for the rest as weighted sums of the numbers of mismatches in the seed region, the numbers of mismatches in the non-seed regions, and an extra cost factor for secondary PAMs. A CasValue exclusion parameter –x defines the upper limit of scores of footprints that will be considered potential Cas9 binding sites for a candidate site’s targeting sequence, so that only footprints with scores ≤ –x are counted as possible off-targets and the rest ignored. This count is compared with an ‘accept’ number specified with the candidate site’s CasFinder input sequence, and those candidates whose counts are ≤ the ‘accept’ number are accepted by CasFinder as specific Cas9 targets and reported in its output file. All scoring parameters are under user control but default to 1 for each mismatch (for both seed and non-seed), 1 for the secondary PAM cost, and 3 for –x. When CasFinder is used to search for targets endogenous to the genome (see item 4 below) the ‘accept’ number defaults to 1; otherwise the default is 0. These ‘accept’ defaults are appropriate for typical usages, as a footprint corresponding to a Cas9 target site with a score within the –x threshold will always be found if the target site is endogenous to the genome (the footprint being the target site itself), but should not be found if the Cas9 target site comes from sequence exogenous to the genome. However, occasions arise where it is useful to override these defaults. For instance, accept numbers > 1 can be used to locate endogenous Cas9 target sites in genes for which multiple copies exist in the genome. A benefit of the CasValue scoring system is that for any accepted candidate Cas9 target, CasValue identifies and reports the lowest score of any Cas9 footprint found in the genome that is > –x as a “threshold rejection score” (TRS). In broad terms, the TRS indicates the distance in sequence space to the closest potential Cas9 off-target site in the genome outside of any of its accepted instances. It can thus be used to rank accepted Cas9 sites by specificity, with higher TRSes indicating more specificity.
3. To make it easy to add new Cas9s and genomes to the system, or adjust Cas9 characteristics such as PAM sequences and seed positions, all information about Cas9s and genomes is maintained in user-editable configuration files so that such modifications never require changes to program code. These are supplemented with a wide array of program and input file options that allow additional fine tuning of scoring and Cas9 parameters in each CasFinder run.
4. The CasFinder system also provides additional usability features, such as the ability to specify a ‘reference point’ within each input sequence to be searched for Cas9 targets. CasFinder reports the offsets of any accepted Cas9 targets relative to these reference points. If, for instance, the reference point is set to the 5’ end of an exon in the sequence, these offsets make it easy to identify targets in the sequence close to the beginning of the exon. Additionally, when users desire to search for endogenous targets in genome regions of interest, users may specify these in the CasFinder input file just by providing FASTA headers containing the regions’ location ranges instead of providing full FASTA records complete with nucleotide sequence. CasFinder will extract these regions itself from the genome sequences declared in its configuration files.

The CasFinder system has been implemented as a pair of perl programs CasValue and CasFinder, a pair of text configuration files, and two perl packages that interface with the configuration files. To install the software requires editing the configuration files and, if necessary, generating bowtie indexes for genomes to be processed. Detailed instructions on installation and usage of the system are in Supporting Information S1. The CasFinder system has been tested and run on Debian GNU/Linux 6.0 for amd64 CPUs. The system was developed and run using perl 5.10.1 but has been tested and found to generate identical output on perl 5.18.1.

CasFinder system performance depends heavily on the number of matches returned by its bowtie queries. Given the use of queries accommodating seed mismatches, these numbers can be very large, especially in large genomes. For this reason, CasFinder is set to ignore candidate targets overlapping repeatmasked sequence by default, and the seed sequence lengths specified in distributed configuration files are set at 13bp, larger than the 11-12bp usually considered. However, even sequences containing no repeatmasked bases can return large numbers of matches: Indeed, in testing we encountered an exonic seed query containing no repeatmasked bases that returned 692180 human genome matches. By default CasValue has bowtie return all matches to a query, but to allow users to control performance an option is provided pass bowtie a –k parameter [25] which limits the maximum number of bowtie returns per query to the specified value. When bowtie –k is used and a query for a candidate target seed returns exactly –k matches, CasValue cannot be certain it has evaluated all the candidate’s bowtie-retrievable Cas9 footprints in the genome. CasValue will still accept such candidates if the candidate’s ‘accept’ limit has not been exceeded, but it will report them as incompletely analyzed. Compared to evaluation without a –k parameter, such sites might be incorrectly reported in either of two ways: They might have been rejected instead of accepted if more footprints had been examined, or they might have been accepted but have lower TRSes. A test with 30000 randomly picked exonic repeatmasked SP Cas9 targets in the hg19 human genome found that a roughly constant fraction of between 19 and 34% (mean 28%) of incompletely analyzed targets were both correctly accepted and had correct TRSes at a range of –k limits between 1000 and 20000 (see Supporting Information S1, Figure S5). Users who specify a –k value and are concerned with incompletely analyzed sites can re-evaluate these sites at higher –k or have bowtie return all matches.

## Results

We used the CasFinder system to generate a catalog of Cas9 sites across the human and mouse exomes for the SP, ST1, and NM Cas9s. Configuration file parameters for the Cas9s are given in Table 1, and statistics on the output and performance of the runs are provided in Table 2. Files of all Cas9 sites are available on our web site (http://arep.med.harvard.edu/CasFinder). To our knowledge this is the first attempt to find Cas9 sites on an exome-wide scale for Cas9s other than SP.

**Table 1:** Cas9 configuration parameters for analysis of human and mouse exomes. ST1 and NM PAMs are taken from ref.[11].

| Cas9 | Target length | Seed position |  | Primary PAM(s) | Secondary PAM(s) |
| --- | --- | --- | --- | --- | --- |
|  |  | Start | end |  |  |
| SP | 20 | 8 | 20 | NGG | NAG |
| ST1 | 20 | 8 | 20 | NNRGAA,NNAGGA | NNANAA,NNGGGA |
| NM | 20 | 8 | 20 | NNNNGANN,NNNNGTTN,NNNNGNNT | NNNNGTNN,NNNNGNTN |

**Table 2:**
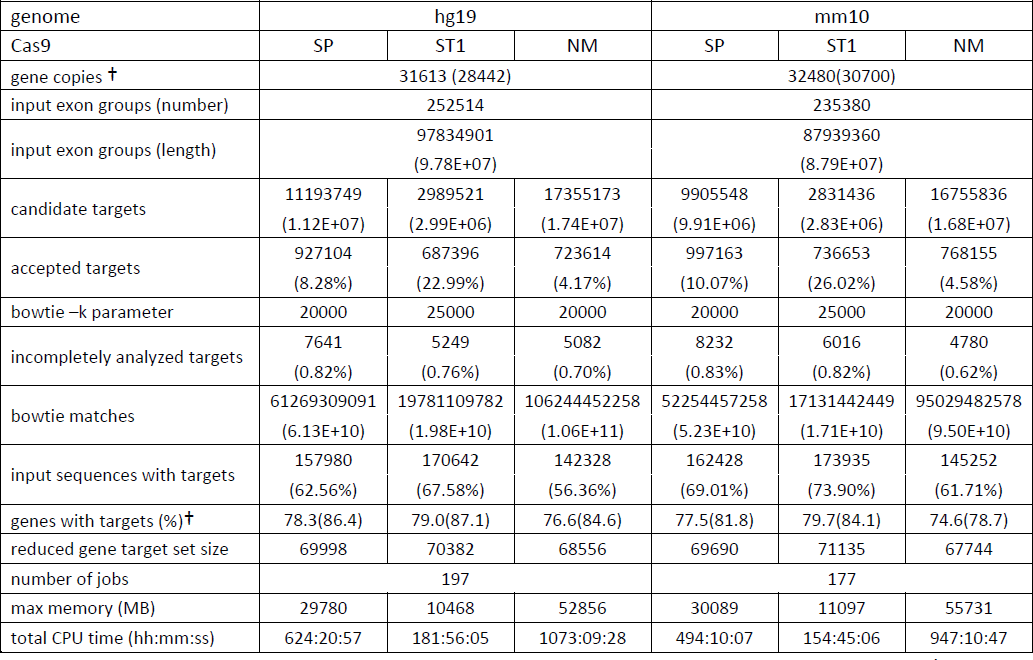
CasFinder and CasValue output and performance when applied to the human and mouse exomes. † Values in parentheses do not distinguish multiple copies of genes in the genome (see Supporting Information S1).

The human and mouse genomes and exomes were derived from chromosome sequences and exon data downloaded from the UCSC Genome Browser (version GRCh37/hg19 for human and GRCm38/mm10 for mouse) [26,27]. Exonic sequence ranges were assigned to each UCSC-recognized geneSymbol by merging the ranges of all exons from all isoform annotations associated with the geneSymbol, where geneSymbols mapped to multiple regions of the genome (multi-copy genes) were first disambiguated. The merged exon sequence ranges are designated as *exon groups* (see Table 2) that are then numbered in 5’-to-3’ order for each geneSymbol. Because the same exon may have a different numbers in different isoforms due to alternative splicing and transcription starts, exon group number *n* may not correspond to exon *n* in many (or even any) originating isoforms. Exon group sequence ranges were padded by 20bp at each end to facilitate finding sites at their edges.

In general, for both human and mouse, the numbers of candidate sites go in an order NM > SP > ST1 that is consistent with the degree of degeneracy of primary PAM motifs (see Table 1). For instance, the ST1 primary PAM specifies three motifs of four fixed bp (breaking the ambiguous R base into a separate A and G), while the SP PAM specifies two fixed bp: If bases were equally distributed, ST1 PAMs would be 18.75% as frequent as SP PAMs, and NM would generate twice as many. However, the order of *accepted* candidates is rearranged to SP > NM > ST1. This is possibly because more degenerate PAMs not only increase the number of candidate sites, but also increase the likelihood that a genomic match of a candidate seed will be followed by a primary or secondary PAM, and thereby the chance that this match will be scored as an off-target that causes rejection of the candidate.

As expected from system design, the number of candidate sites for a Cas9 also correlates with the computer resources required to process it. It required ∼5x as much memory and ∼6x as much CPU time to process NM *vs.* ST1, with SP in between. To process an entire exome given available computer resources, exome regions were divided into parts of up to 500kbp each (197 for human and 177 for mouse), and each part processed as a separate job, where jobs ran in parallel on a computer cluster. In this way, analysis of an entire exome could be performed in less than a day. To further control computer resource requirements, the bowtie –k option was used with values chosen to bring the fraction of incompletely analyzed sites to < 1%. The ST1 Cas9 required a larger –k than either SP or NM (25000 *vs.* 20000).

The whole exome Cas9 site catalogs yielded between 687K and 997K sites and targeted 56-74% of all exons in 77-80% of genes in the genome (see Table 2), and can be used to identify sites targeting critical domains or exons of genes of interest. For convenience in choosing sites from these catalogs, BED files were generated for each catalog that can be loaded as custom tracks into the UCSC Genome Browser [27,28], allowing users to visualize where the sites are relative to their targeted exons, genes, and other genomic features. Some genes or exons of interest may not be represented in the catalogs: Common reasons for this include the presence of repetitive sequence, or the presence of homologs or pseudogene copies in the genome. Users interested in targeting such genes or exons could rerun CasFinder in these regions with options to process repeatmasked sequence in the former case, or that increase the ‘accept’ number in the latter; decreasing the –x parameter could also be helpful. For use of Cas9 in high-throughput gene knockout screens using pools of sgRNAs [22-24], it is desirable to have reduced catalogs of sites that target the 5’ regions of genes. We generated reduced catalogs of up to three 5’-targeting Cas9 sites per gene for each genome and each Cas9, each containing 68.5K-71.1K sites (see Table 2). Targets in the reduced catalogs were chosen by picking the 5’-most site in each of the first three exon groups for which sites were found, picking more sites in earlier exon groups if less than three site-containing groups existed (see Supporting Information S1). This heuristic was meant to strike a balance between picking sites closest to the 5’ end of the gene, which have most potential to generate a disrupting mutation, while hedging against the possibility that the 5’-most exon might not be expressed in a given cell type. More than 98% of genes with targets in the full exome catalogs are represented in the reduced sets (see Supporting Information S1, Table S7).

### Usage considerations

Although the CasFinder system’s identification and scoring of specific Cas9 target sites results from a systematic and comprehensive calculation, its evaluation is ultimately heuristic. CasValue’s scoring metric only approximately represents factors that have been shown to bear on Cas9 specificity, and its search through the genome is limited to examination of Cas9 footprints with up to a fixed number of mismatches from a candidate target’s seed sequence. The assumption of a fixed length seed sequence is also an idealization and, to improve performance of large scale target searches such as the exome-wide searches performed here, the seed is set in the configuration files distributed with this article to 13bp, larger than often assumed in other studies. However, heuristics cannot be avoided given the current state of knowledge of Cas9 specificity. Currently available data are insufficient to do more than provisionally estimate the sensitivity and accuracy of CasFinder and other Cas9 site evaluation algorithms for finding sites without off-targets; however, we provide a preliminary assessment and discussion of CasFinder sensitivity, specificity, and false discovery rate in Supporting Information (S1). The estimates suggest that CasFinder is a relatively conservative algorithm. Finally, specificity must be generally balanced against the ability to find target sites, as specificity constraints that are too stringent will cause all candidate sites in a region of interest to be rejected. The CasFinder system is meant to provide a way of balancing these needs that is both intuitive and easy to adjust to circumstances. The system’s default settings will result in finding sites for which the targeting sgRNA will have at least 2 mismatches in the seed region of any PAM-adjacent sequences in the genome, and 4 mismatches with any other PAM-adjacent sequences (or 3 mismatches with a secondary PAM). We have found that 4bp of sequence divergence is generally enough to reduce Cas9 activity to background levels [12] (although exceptions have been reported [17]). Targets generated using the CasFinder system are now in use in our laboratory. Higher specificity, if needed, can be requested by increasing the –x parameter, and lower specificity by decreasing –x or by increasing the ‘accept’ number of the target, and scoring sensitivity to Cas9 target site features can also be modulated at a more detailed level by changing scoring weights or seed handling. Finally, seed lengths and seed mismatch limits can be overridden to ensure increased scrutiny of matching seed regions in the genome, although we recommend that this be done selectively and by using CasValue to re-screen targets found initially through CasFinder searches with default parameters (see Supporting Information S1, Table S5).

The CasFinder system takes its place among a heterogeneous set of reported Cas9 target analysis algorithms and software tools, reflecting the broad range of applications being developed for CRISPR technology, each with its own specific priorities and needs. A comparison of the CasFinder system with four recently reported similar methods, including the CRIPSR Design Tool [18] and CasOT [29], and two recent studies that generated human exome-wide Cas9 target sets [22,23], is provided in Supporting Information S1 (see Table S8). This comparison shows that there are now a wide variety of approaches and priorities in evaluating Cas9 targets, but also suggests that there is need at a software level for tools that enable users to flexibly tune and tweak the selection of Cas9 targets to the needs of their research. The CasFinder system is offered as an initial entry into this niche that is both immediately useful and that, we hope, can serve as an example of a software architecture that increases user control over Cas9 analysis algorithms that other research groups can adopt or further improve.

## Availability and future directions

The perl code, example configuration files, and instructions are all provided on our supplemental web site http://arep.med.harvard.edu/CasFinder under the OSI-compliant MIT license, along the whole exome and reduced catalogs of Cas9 sites generated for the analyses in Table 2. We anticipate incorporating new scoring metrics and generating large numbers of genome-wide target sets for a multiplicity of species as community resources, as well as generating target sets with new Cas9s that are developed for genome editing [30]. We call on the research community to provide feedback on developing additional algorithms and functionality for Cas9 targeting software.

## Acknowledgements

This work was supported by NIH grant P50 HG005550. We thank Amir Karger for assistance using Harvard Medical School’s Orchestra High Performance Compute Cluster, both supported by NIH grant NCRR 1S10RR028832-01.

## Supporting Information

Figure S1. Flow chart of high level logic of CasFinder program.

Figure S2. Flow chart of high level logic of CasValue program.

Text S1. CasFinder System Documentation: Strategy, Installation, Usage, and Performance; also contains Tables S1-S8 and Figures S3-S5.

